# Reconstruction and Simulation of a Scaffold Model of the Cerebellar Network

**DOI:** 10.1101/532515

**Authors:** Stefano Casali, Elisa Marenzi, Chaitanya Medini, Claudia Casellato, Egidio D’Angelo

## Abstract

Reconstructing neuronal microcircuits through computational models is fundamental to simulate local neuronal dynamics. Here a *scaffold model* of the cerebellum has been developed in order to flexibly place neurons in space, connect them synaptically and endow neurons and synapses with biologically-grounded mechanisms. The scaffold model can keep neuronal morphology separated from network connectivity, which can in turn be obtained from convergence/divergence ratios and axonal/dendritic field 3D geometries. We first tested the scaffold on the cerebellar microcircuit, which presents a challenging 3D organization, at the same time providing appropriate datasets to validate emerging network behaviors. The scaffold was designed to integrate the cerebellar cortex with deep cerebellar nuclei (DCN), including different neuronal types: Golgi cells, granule cells, Purkinje cells, stellate cells, basket cells and DCN principal cells. Mossy fiber (mf) inputs were conveyed through the glomeruli. An anisotropic volume (0.077 mm^3^) of mouse cerebellum was reconstructed, in which point-neuron models were tuned toward the specific discharge properties of neurons and were connected by exponentially decaying excitatory and inhibitory synapses. Simulations using pyNEST and pyNEURON showed the emergence of organized spatio-temporal patterns of neuronal activity similar to those revealed experimentally in response to background noise and burst stimulation of mossy fiber bundles. Different configurations of granular and molecular layer connectivity consistently modified neuronal activation patterns, revealing the importance of structural constraints for cerebellar network functioning. The scaffold provided thus an effective workflow accounting for the complex architecture of the cerebellar network. In principle, the scaffold can incorporate cellular mechanisms at multiple levels of detail and be tuned to test different structural and functional hypotheses. A future implementation using detailed 3D multi-compartment neuron models and dynamic synapses will be needed to investigate the impact of single neuron properties on network computation.

## 1 Introduction

The causal relationship between components of the nervous system at different spatio-temporal scales, from subcellular mechanisms to behavior, still needs to be disclosed and this represents one of the main challenges of modern neuroscience. The issue can be faced using *bottom-up* modeling, which allows propagating microscopic phenomena into large-scale networks (Markram, 2012; Markram et al., 2015; D’Angelo and Wheeler-Kingshott, 2017). This *reverse engineering* approach integrates available details about neuronal properties and synaptic connectivity into *realistic* computational models and allows to monitor the impact of microscopic variables on the integrated system. Realistic modeling can predict emerging collective behaviors producing testable hypotheses for experimental and theoretical investigations (Llinas, 2014) and might also play a critical role in understanding neurological disorders (Soltesz and Staley, 2018). In practice, realistic modeling of microcircuit dynamics and causal relationships among multi-scale mechanisms poses complex computational problems. First, the modeling strategy needs to be flexible accounting for a variety of neuronal features and network architectures, to be easy to update as new anatomical or neurophysiological data become available, and to be easy to modify in order to test different structural and functional hypotheses. Secondly, the modeling tools need to be scalable to the dimension of the network and to the nature of the scientific question (Destexhe et al., 1996), to be suitable for available simulation platforms, e.g. pyNEST and pyNEURON (Brette et al., 2007; Hines et al., 2009), and to efficiently exploit High-Performance Computing (HPC) resources.

Markram and colleagues recently carried out a digital reconstruction of the neocortical microcolumn by integrating experimental measurements of neuronal morphologies, layer heights, neuronal densities, ratios of excitatory to inhibitory neurons, morphological and electro-morphological composition, electrophysiological properties of neurons and synapses (Markram et al., 2015). Neuron parameters were derived from databases specifically addressing cerebro-cortical neuron properties (e.g. Blue Brain Project and Allen Brain Atlas (Markram, 2006; Sunkin et al., 2013)). Microscopic network wiring was then estimated computationally through a touch detection algorithm, that is based on a probability/proximity rule (i.e. the probability that morphologically defined dendrites and axons make a synaptic connection depends on their spatial proximity). This approach, in which the reconstruction of microcircuit connectivity depends on the 3D morphology of the axonal and dendritic processes of individual cells, may apply to brain structures for which datasets comparable to neocortex are available. However, such specific datasets are not available in general for all brain regions and it seems convenient in principle to keep separated neuronal morphology from network connectivity, which is reported as convergence/divergence ratios and axonal/dendritic field geometries in the literature in many cases.

The cerebellum hosts the second largest cortical structure of the brain and contains about half of all brain neurons. Modeling the cerebellum brings about specific issues reflecting the peculiar properties of this circuit, which shows a quasi-crystalline geometrical organization well defined by convergence/divergence ratios of neuronal connections and by the anisotropic 3D orientation of dendritic and axonal processes (Fig. 1)(D’Angelo et al., 2016).Moreover, the morphological reconstruction of axonal and dendritic processes of cerebellar neurons is not as developed as for other brain microcircuits, like cerebral cortex and hippocampus (e.g. see the NeuroMorpho.org repository – https://www.re3data.org/repository/r3d100010107) (Akram et al., 2018). Therefore modeling the cerebellum relies on a knowledge base that differs from that available for the cerebral cortex and thus requires a more general approach than in the Markram’s modeling workflow (Markram et al., 2015).

**Fig. 1.**
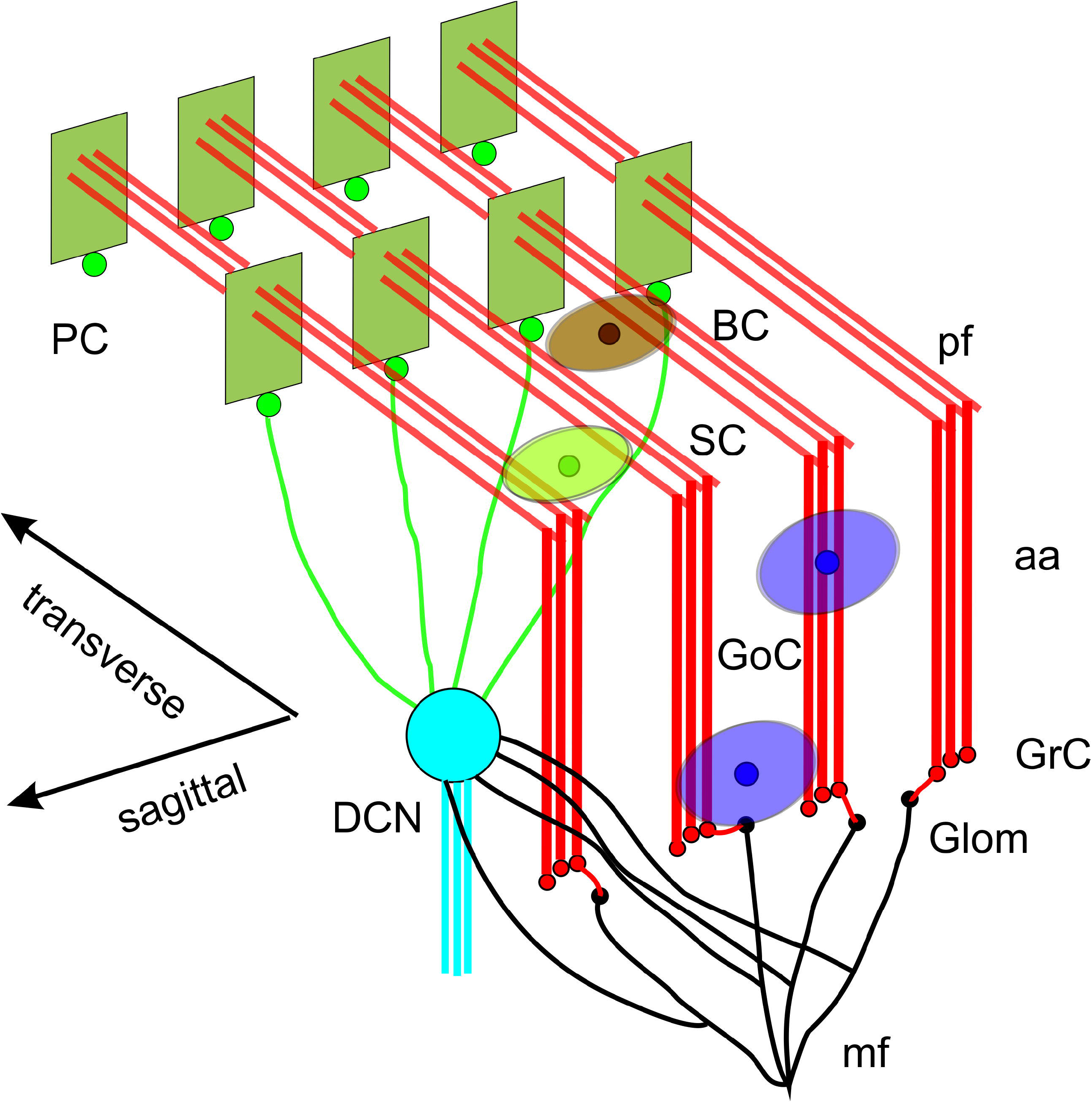
Reconstruction of a scaffold model of the cerebellar network. Schematic representation of the cerebellar network (from (D’Angelo et al., 2016)). Glomeruli (Glom); mossy fiber (mf); Granule cells (GrC); ascending axon (aa); parallel fiber (pf); Golgi cells (GoC); stellate cell (SC); basket cell (BC); Purkinje cell (PC); Deep Cerebellar Nuclei cell (DCNC). Gloms transmit mf inputs to GrCs, which emit aa and pf, which in turn activate GoCs, PCs, SCs and BCs. GoCs inhibit GrCs, SCs and BCs inhibit PCs. DCN cells are inhibited by PCs and activated by mf. Note the precise organization of PC dendrites, SC/BC dendrites and GoC dendritic arborization on the parasagittal plane. The same abbreviations are used also in the following figures.

Some recent models were purposefully designed to reproduce a limited section of the cerebellar cortex, the granular layer (Maex and De Schutter, 1998; Solinas et al., 2010; Sudhakar et al., 2017), in great detail and incorporated Hodgkin-Huxley-style mechanisms and neurotransmission dynamics (D’Angelo et al., 2001; Solinas et al., 2007a, b; Nieus et al., 2014; Masoli et al., 2015; Masoli and D’Angelo, 2017; Masoli et al., 2017). Other models were designed to simulate, in a simplified form, large-scale computationally efficient networks of the olivo-cerebellar system (Medina and Mauk, 2000; Yamazaki and Nagao, 2012). In this work, a new cerebellum *scaffold model* has been developed and tested, allowing to incorporate axonal/dendritic field geometries specifically oriented in a 3D space and to reconnect neurons according to convergence/divergence ratios typically well defined for the cerebellum (D’Angelo et al., 2016). The cerebellum scaffold model maintains scalability and can be flexibly handled to incorporate neuronal properties on multiple scales of complexity and to change its connectivity rules. For the sake of simplicity, here we used first simplified neuron and synaptic models to evaluate the impact of construction rules. The cerebellum scaffold model was validated by testing its ability to reproduce the structural properties anticipated experimentally and the emergence of complex spatiotemporal patterns in network activity. The model was run on the pyNEST and pyNEURON simulators (Brette et al., 2007; Hines et al., 2009) and a test workflow was integrated into a large-scale neuroinformatics infrastructure, the Brain Simulation Platform (https://collab.humanbrainproject.eu/).

## 2 Materials and Methods

This paper reports the design and implementation of a *scaffold* model of the cerebellum microcircuit. The model architecture is scalable and is designed to host different types of neuronal models and to determine their synaptic connectivity from convergence/divergence ratios and axonal/dendritic field geometries reported in literature. The workflow encompasses two main modules in cascade: **cell placement** into a user-defined volume; **connectivity** among neurons. The scaffold can then be used for **functional simulations** of network dynamics (Fig. 1). The scaffold is designed to be embedded into different simulators, e.g. pyNEST and pyNEURON. This workflow, by allowing a flexible reconstruction of the cerebellar network, will eventually allow to evaluate physiological and pathological hypotheses about circuit functioning.

### 2.1 Cell Placement Module

The *Cell Placement Module* places the cells in a virtual network volume divided in layers based on morphological definitions. The process takes into consideration the different cerebellar neuron types: the Golgi Cell (GoC), Granule Cell (GrC), Purkinje Cell (PC), Stellate cell (SC),Basket Cell (BC), Deep Cerebellar Nuclei glutamatergic GAD-positive cell (DCNC), and glomerulus (Glom). Glom is actually a mossy fiber terminal and is represented as a neuronal element at the input stage, while DCNCs are placed at the output stage of the circuit. For each neuron type, the density value in a specific layer was derived from literature, and geometric features (including soma radius and 3D-oriented dendritic and axonal fields) were defined according to experimental data. Through *ad hoc* algorithms (*Bounded Self-Avoiding Random Walk Algorithm* and *Purkinje Cells placement algorithm*, see below), the cells were positioned in the 3D volume of each layer, according to their density, soma radius, and anisotropic extension, ensuring that their somata did not overlap. The module was implemented in Python, and its output was saved in an *.hdf5* file containing the unique identification number (ID) of each cell, its corresponding type (an integer value between 1 and 7, as in Table 1), and the three spatial coordinates of the soma center (x, y, z). To evaluate the effectiveness of the cell positioning algorithms, we derived a continuous distribution of pair-wise distances using *kernel density estimation* (KDE), in which the Gaussian kernel had fixed bandwidth for each cell population. KDE yielded a single maximum when pair-wise distances were distributed homogeneously (GrC, GoC, SC, BC, DCNC) and multiple local maxima when distances were placed according to different geometric rules (PC). A reconstructed network volume and pair-wise soma distances yielded by this module are illustrated in Fig. 2.

**Fig. 2.**
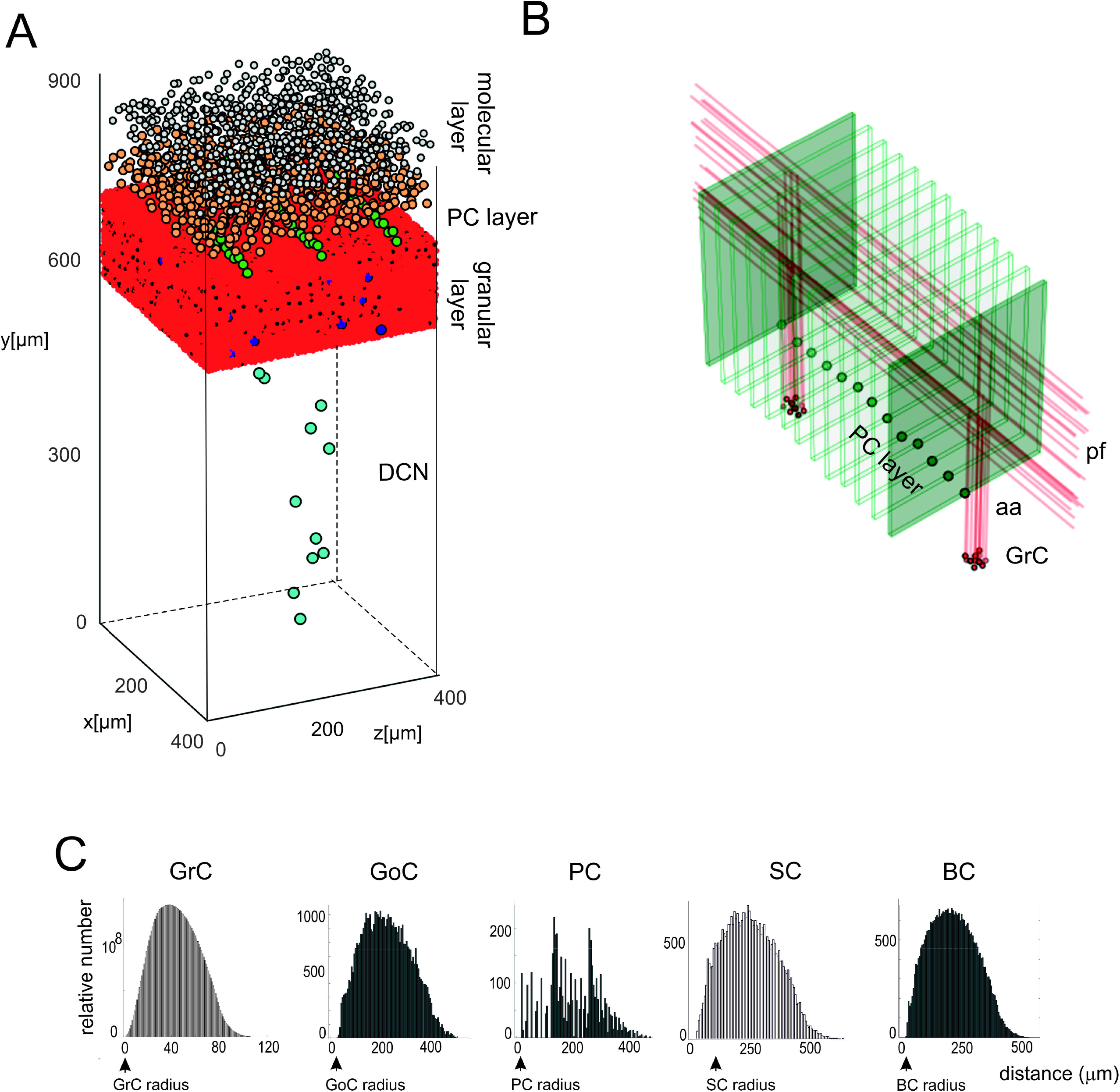
Cell placement and network architecture. (**A**) The cells are placed in the network 3D space using a Bounded Self-Avoiding Random Walk Algorithm. The figure shows the volume of 400 x 400 x 900 μm^3^ containing 96.737 neurons and 4.220.752 synapses used for simulations. (**B**) Projection of GrC axons to the molecular layer hosting the PCs (green dots in the PC layer are the somata, the thin green parallelepipeds above are the corresponding dendritic trees occupying the molecular layer). The figure shows two clusters of GrCs and the corresponding aa and pf, illustrating that the cerebellar network connectivity respects the 3D architecture shown in Fig. 1. (**C**) Distributions of 3D pair-wise inter-soma distances within each neuronal population: GrCs, SCs, GoCs, BCs and PCs. Note that the distributions are nearly normal, except for the PCs.

**Table 1.**
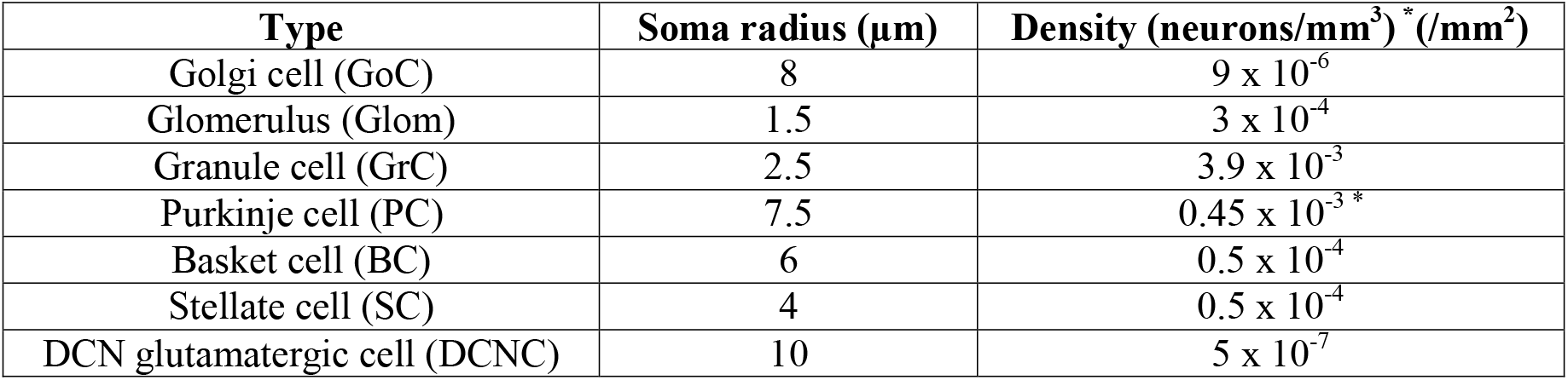
Neuron types, size and density. The table reports the density of neurons in the layer volume (neurons/mm^3^), except for PCs for which the planar density is used (neurons/mm^2^).Data for Glom, GrC, GoC, PC, SC, BC from (Korbo et al., 1993). The density of DCNC was estimated from the ratio of PCs to DCNCs(11:1)(Person and Raman, 2012).

| Type | Soma radius (μm) | Density (neurons/mm<sup>3</sup>) * (/mm<sup>2</sup>) |
| --- | --- | --- |
| Golgi cell (GoC) | 8 | $9 \times 10^{-6}$ |
| Glomerulus (Glom) | 1.5 | $3 \times 10^{-4}$ |
| Granule cell (GrC) | 2.5 | $3.9 \times 10^{-3}$ |
| Purkinje cell (PC) | 7.5 | $0.45 \times 10^{-3}$* |
| Basket cell (BC) | 6 | $0.5 \times 10^{-4}$ |
| Stellate cell (SC) | 4 | $0.5 \times 10^{-4}$ |
| DCN glutamatergic cell (DCNC) | 10 | $5 \times 10^{-7}$ |

GrCs, GoCs, SCs, BCs, and DCNCs were placed in thin sub-layers (with height = 1.5x soma diameter) using a bounded self-avoiding random walk algorithm. In each sub-layer, the cells were initially distributed in 2D and then sub-layers one on top of the others. The first cell was placed randomly and each subsequent one was positioned nearby along a random angular direction. The overlap among somata was avoided since, along the selected direction, the minimum distance to place the next cell was the soma diameter. A random term was added to the minimum distance to scatter the somata (the range depended on neuron density). If the surrounding space was completely occupied, i.e. there was insufficient space to place a further cell, a new starting point was selected resetting the random walk process for the remaining neurons in that sub-layer. Once completed, the 2D sub-layer was piled on top of the underlying one and a random noise was added to soma position on the vertical coordinate (Korbo et al., 1993). This approach maintained randomness achieving a realistic quasi-Gaussian distribution of pair-wise inter-neuron distances (see Fig. 2C) and proved computationally efficient.

The PCs were distributed in a single sub-layer forming an almost planar grid between the granular and molecular layers. The PC inter-soma distances over this plane were constrained by the dendritic trees, which are flat and expand vertically on the parasagittal plane (about 150 μm radius x 30 μm width) without overlapping (Masoli et al., 2015). Since PC somata do not arrange in parallel arrays but are somehow scattered, a noisy offset was introduced creating an average angular shift of about 5° between adjacent PCs. As for the other neuron types, a small random noise was also imposed on the vertical coordinate (Korbo et al., 1993).

The data required for cell positioning in the cerebellar cortex were obtained from literature (Eccles et al., 1967; Magyar et al., 1972; Mezey et al., 1977; Hamori and Somogyi, 1983; Jakab and Hamori, 1988; Korbo et al., 1993; Sultan, 2001; Santamaria et al., 2007; Barmack and Yakhnitsa, 2008; Solinas et al., 2010) and are summarized in Table 1. GoCs, GrCs and Gloms were placed into the granular layer; BCs and SCs in the lower and upper half of the molecular layer, respectively. A certain number of DCNCs was randomly distributed in DCN volume according to the PC/DCNC ratio, since more specific parameters are still missing (Gauck and Jaeger, 2000; Aizenman et al., 2003; Person and Raman, 2012). Special care was given to the GrC ascending axon (aa) that, starting directly from the soma, makes its way up vertically toward the molecular layer. The height of each ascending axon was chosen from a Gaussian distribution in the range of 181±66 μm (Huang et al., 2006). This value represents also the vertical coordinate of the corresponding parallel fiber (pf), running transversally and parallel to the cerebellar surface.

### 2.2 Connectivity Module

The connectivity module created structural connections between pairs of neurons belonging to specific types. Each neuron type formed input and output connections with other neurons of the same or different types. Therefore, once the placement was completed, it was possible to reconstruct the connectome applying connectivity rules based on *proximity* of neuronal processes and on *statistical ratios* of convergence and divergence. When available, morphological and statistical literature data were used, otherwise plausible physiological constraints were applied. In our scaffold, 16 connection types were generated (the most important are shown in Fig. 3), from the volume covered by pre-synaptic axonal processes to that covered by post-synaptic dendritic trees of specific neuron types:

1. From glomeruli to granule cells(Glom-GrC);
2. From glomeruli to basolateral dendrites of Golgi cells (Glom-GoC);
3. From Golgi cells to Gloms (GoC-Glom): this is fused together with Glom-GrC to generate directly GoC-GrC connections;
4. Among Golgi cells (GoC-GoC);
5. From ascending axons of granule cells to Golgi cells (aa-GoC);
6. From parallel fibers of granule cells to apical dendrites of Golgi cells (pf-GoC);
7. Among stellate cells (SC-SC);
8. Among basket cells (BC-BC);
9. From parallel fibers of granule cells to stellate cells (pf-SC);
10. From parallel fibers of granule cells to basket cells (pf-BC);
11. From stellate cells to Purkinje cells (SC-PC);
12. From basket cells to Purkinje cells (BC-PC);
13. From ascending axons of granule cells to Purkinje cells (aa-PC);
14. From parallel fibers of granule cells to Purkinje cells (pf-PC);
15. From Purkinje cells to DCN cells (PC-DCNC);
16. From glomeruli to DCN cells (Glom-DCNC).

**Fig. 3.**
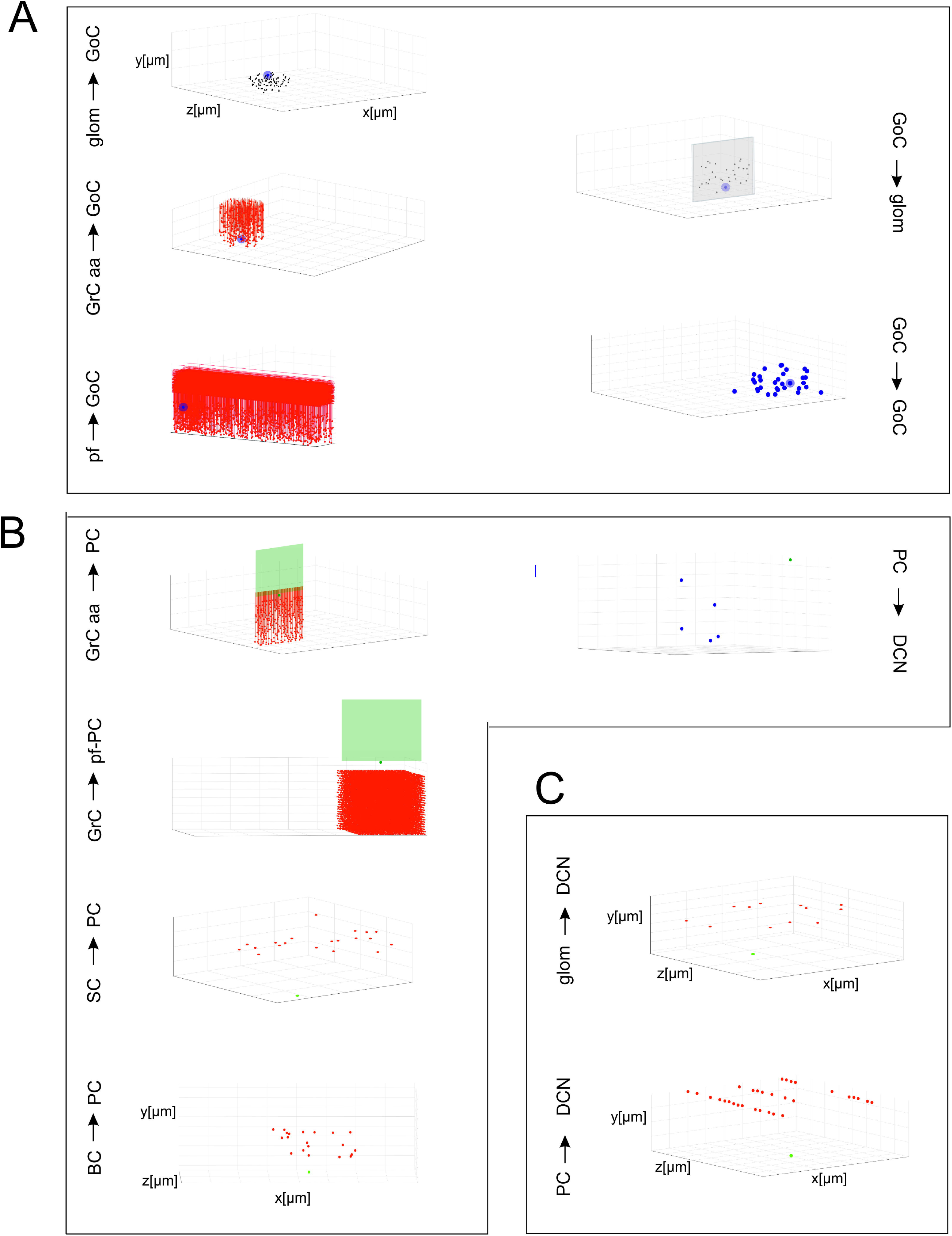
Cell connectivity: examples for specific connections. Examples of divergence and convergence at different connections in the cerebellar network space. The plots have base area (400×400 μm^2^) and thickness specific for each layer. The plots show a randomly selected pre-synaptic cell together with its connected post-synaptic neurons (divergence) or viceversa (convergence). (**A**) Connections of GrCs and GoCs. (**B**) Connections of PCs, SCs and BCs. (**C**) Connections of DCNCs.

Given a connection type, for each pre-synaptic neuron, the potential post-synaptic cells were identified as those that met geometric neuron-specific constraints. Then, given the convergence and divergence ratios, post-synaptic neurons were selected among the potential ones, through a pruning process proceeding through distance-based probability functions specific for each volume direction. The module was implemented in Python, and its output saved in an *.hdf5* file containing a matrix for each connection type, in which each row was defined by three values: the unique ID of the pre-synaptic neuron, the unique ID of the post-synaptic neuron and the inter-soma 3D distance between that pair (see Fig. 4A). The plots in Fig. 4B-Ccompare, for each connection type, the divergence and convergence ratios reported by literature to the values obtained after scaffold reconstruction in a sample volume. The cell placement and connections rules yielded indeed a very good approximation of the anatomical and physiological parameters reported in literature.

**Fig. 4.**
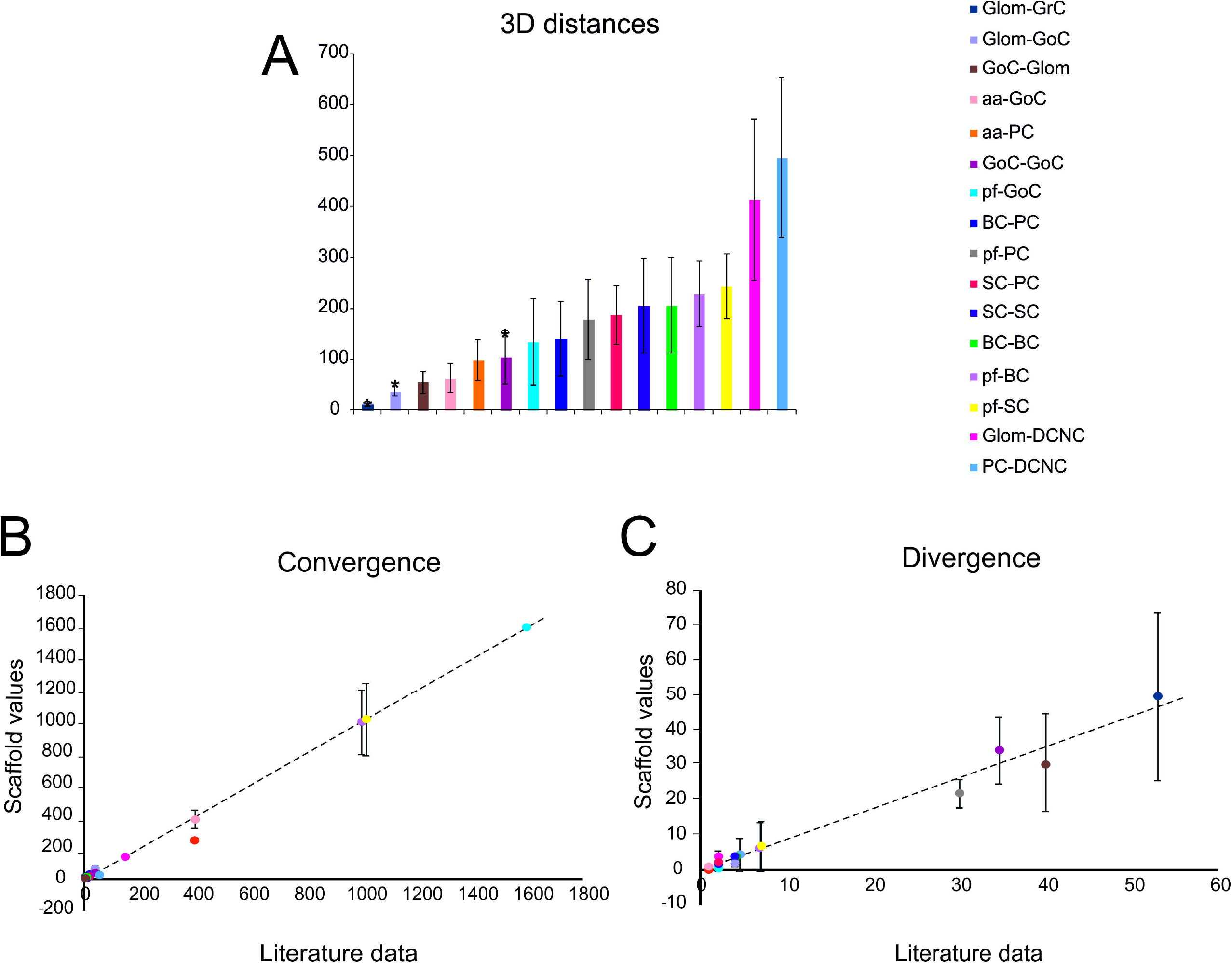
Cell connectivity: pair-wise distance prediction and convergence/divergence validation. (**A**) Pair-wise distance prediction deriving from the placement and subsequent cell-to-cell connectivity. The data that find correspondence in literature are indicated as asterisks. For each connection type, the pair-wise distances between connected cells (inter-soma distance) are reported. Data from: (1) (D’Angelo et al., 2013), (2) (Barmack and Yakhnitsa, 2008), (3) (Rieubland et al., 2014). (**B-C**) The plots compare divergence and convergence for the different connections of the scaffold with those anticipated experimentally. The regression lines show a very close correspondence of the model to experimental results. Linear regression lines are fitted to the data (divergence: r^2^ = 0.98, slope = 0.88; convergence: r^2^ = 0.99, slope = 0.99). Data from: (1) (Nieus et al., 2006), (2) (Dieudonne, 1998), (3) (D’Angelo et al., 2013), (4)(Solinas et al., 2010), (5) (Kanichay and Silver, 2008), (6) (Cesana et al., 2013), (7) (Hull and Regehr, 2012), (8) (Lennon et al., 2014), (9) (Huang et al., 2006), (10) (Jorntell et al., 2010), (11) (Sultan and Heck, 2003), (12) (Person and Raman, 2012), (13) (Boele et al., 2013).

### 2.3 Functional Simulations

In order to test the functionality of the scaffold, single neuron models were placed in the corresponding positions of the connectome matrix. In this first version of the cerebellar microcircuit, spiking point-neuron models based on Integrate&Fire (I&F) dynamics with conductance-based exponential synapses (i.e. synaptic inputs cause an exponential-shaped change in synaptic conductances) were used. The output files of these simulations contained all the spike events (neuron IDs and relative spike times). Glomeruli were represented as “parrot neurons” just able to pass the imposed stimulation patterns unaltered. Each other neuron type was characterized by specific values, directly related to neurophysiological quantities (*C_m_, τ_m_, E_L_, Δt_ref_, I_e_, V_r_, V_th_*), corresponding to biological values taken from literature available from animal experiments or databases (https://neuroelectro.org/) (Tripathy et al., 2014) (Table 2). In order to account for the neuron-specific dynamics of GABA and AMPA receptor currents, also the decay times of the excitatory and inhibitory synaptic exponential functions (*τ_exc_, τ_inh_*) were set differently for each neuron type (Table 2). Each synaptic connection type was characterized by specific values of weight and delay (Table 3).

**Table 2.**
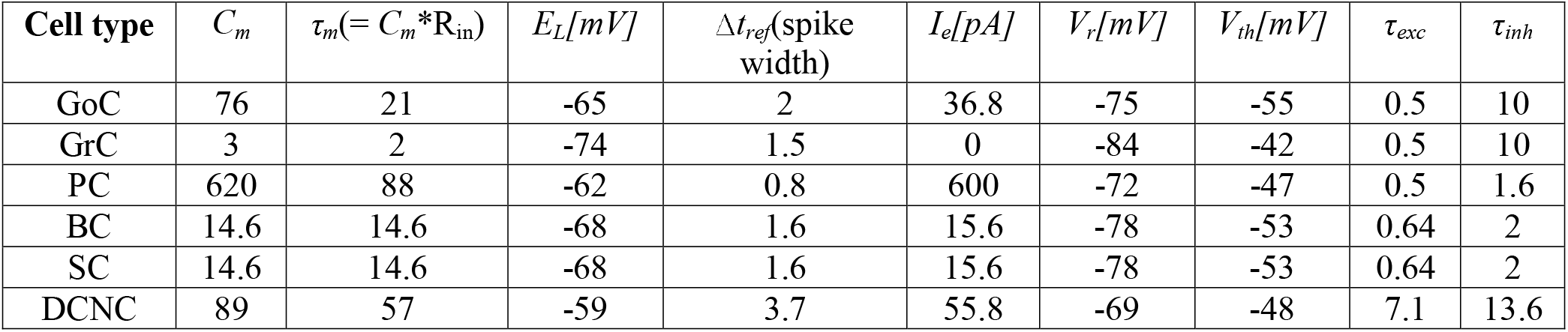
Neuron-specific parameters. The table shows the parameters used to define specific neuronal properties in the model.C_m:_ membrane capacitance; τ_m_: membrane time constant; R_in_: input membrane resistance; E_L_: leakage resting potential; Δt_ref_: refractory period; I_e_: endogenous current; V_r_: reset potential; V_th_: threshold potential; τ_exc_, τ_inh_: excitatory and inhibitory synaptic exponential time constants).Data are obtained from *NeuroElectro* (https://neuroelectro.org/)(Tripathy et al., 2014).

| Cell type | $C_m$ | $\tau_m (= C_m * R_{in})$ | $E_L [mV]$ | $\Delta t_{ref}$ (spike width) | $I_e [pA]$ | $V_r [mV]$ | $V_{th} [mV]$ | $\tau_{exc}$ | $\tau_{inh}$ |
| --- | --- | --- | --- | --- | --- | --- | --- | --- | --- |
| GoC | 76 | 21 | -65 | 2 | 36.8 | -75 | -55 | 0.5 | 10 |
| GrC | 3 | 2 | -74 | 1.5 | 0 | -84 | -42 | 0.5 | 10 |
| PC | 620 | 88 | -62 | 0.8 | 600 | -72 | -47 | 0.5 | 1.6 |
| BC | 14.6 | 14.6 | -68 | 1.6 | 15.6 | -78 | -53 | 0.64 | 2 |
| SC | 14.6 | 14.6 | -68 | 1.6 | 15.6 | -78 | -53 | 0.64 | 2 |
| DCNC | 89 | 57 | -59 | 3.7 | 55.8 | -69 | -48 | 7.1 | 13.6 |

**Table 3.** Synaptic parameters for each connection type. The parameters result from atuning procedure based on data reported in different papers and summarized in (Maex and De Schutter, 1998; Solinas et al., 2010; Sudhakar et al., 2017). A main additional constraint is that the connection weight is larger from aa then pf connections, both for GoCs and PCs (Sims and Hartell, 2005; Cesana et al., 2013).

| Connection types | Weight [nS] | Delay [ms] |
| --- | --- | --- |
| Glom-GrC | 9.0 | 4.0 |
| Glom-GoC | 2.0 | 4.0 |
| GoC-GrC (GoC-Glom-GrC) | -5.0 | 2.0 |
| GoC-GoC | -8.0 | 1.0 |
| aa-GoC | 20.0 | 2.0 |
| pf-GoC | 0.4 | 5.0 |
| SC-SC | -2.0 | 1.0 |
| BC-BC | -2.5 | 1.0 |
| pf-SC | 0.2 | 5.0 |
| pf-BC | 0.2 | 5.0 |
| SC-PC | -8.5 | 5.0 |
| BC-PC | -9.0 | 4.0 |
| aa-PC | 75.0 | 2.0 |
| pf-PC | 0.02 | 5.0 |
| PC-DCNC | -0.0075 | 4.0 |
| Glom-DCNC | 0.006 | 4.0 |

The input stimulus was set by defining the volume where Gloms were activated, the onset time, the duration and the frequency of spikes. A background activity was generated by a Poisson process of stochastic neuronal firing at 1 Hz on all the glomeruli. Superimposed on it, a burst at 150 Hz lasting 50ms (Rancz et al., 2007) was activated 300ms after the beginning of simulation. Indeed, mossy fibers have a low basal activity, but in response to sensorimotor stimuli, can fire at rates beyond 100 Hz. The stimulated volume had a radius of 140μm; the simulation lasted 1 sec, including 300ms pre-stimulus, 50ms stimulus, and 650ms post-stimulus.

In a specific set of simulations(see Fig. 8), we tested the partial contribution of SCs and BCs to the spatiotemporal diffusion of activity among PCs. BCs axonal plexus is preferentially oriented along the parasagittal axis (see (Eccles et al., 1967)). In these simulations, we oriented the SC and BC axonal plexus orthogonally one to each other and concentrated the stimulation burst in a sphere of 30 μm radius.

### 2.4 Network data analysis

For each neuron population, mean frequency rates were extracted in three time windows: baseline pre-stimulus, during stimulus, and steady-state after-stimulus. We then generated peristimulus time histograms (PSTH) for each neuronal population with time bins of 3ms. For each neuron population, we also separated excited from inhibited sub-groups, responding with an increased or decreased firing rate during the stimulus. To do so, we compared the number of spikes during stimulus vs. pre-stimulus normalized by the time-window durations. If the pre-stimulus firing frequency (i.e., baseline) was at least doubled during stimulation, then the cell was classified as excited. Conversely, to classify the inhibited cells. For GrCs, we added a second constraint: to be labeled as excited, a GrC should fire more than 1 spike during stimulation, allowing to exclude spikes determined by the background noise. For DCNC, all cells stopped firing during the stimulation time-window; therefore, no analysis was performed. For each PC, a further *ad-hoc* analysis allowed to identify burst–pause responses. The cells showing a significant stimulus-induced pause (Cao et al., 2012; Herzfeld et al., 2015) were recognized as those in which the first ISI after the end of the stimulus was > 2 SD the pre-stimulus ISI. This comparison was computed within-cell, i.e. for each PC individually.

#### Center-surround analysis

The excitatory-inhibitory balance (*EI*) and center-surround (*CS*) were calculated from firing rates (*FR*) according to (Mapelli and D’Angelo, 2007) and (Solinas et al., 2010) by considering that inhibition occurs only after a delay following the beginning of stimulation. GrC firing rate was then measured 0-20 ms (*T1*) and 20-40 ms (*T2*) after the beginning of stimulation in response to 50 ms at 150 Hz bursts, both in control (*con*) and with GoC-GrC inhibition switched off (*in_off*). The *CS* and *EI* were calculated as follows:

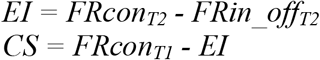

The *CS* values were normalized between 1 and −1. The extension of the center and surround was calculated by including zones with *CS* > 0.5 in the center, and zones with *CS* < −0.5 in the surround. The center and surround relative areas could then be calculated by counting the respective number of pixels and normalizing by the total number of pixels (see Fig. 7C).

#### Oscillation analysis

In order to determine the presence and properties of coherent oscillations in granular layer activity, the activity in a subset of GoCs with overlapping incoming parallel fibers and the related GrCs was analyzed (Maex et al., 2000) during a 5 sec at 5 Hz noisy background mossy fiber activity. The autocorrelations of GoCs and GrCs spike trains and the crosscorrelation between GrCs and GoCs spike trains were calculated using the equation

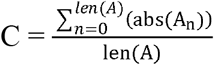

Where C is the index of coherence, A is the array of autocorrelation values, and len(A) is the size of the spike train data array.

The same calculus was executed also for the crosscorrelation

### 2.5 Simulations in pyNEST, pyNEURON and implementation on the Brain Simulation Platform

The microcircuit was implemented and simulated both in pyNEST (Brette et al., 2007; Eppler et al., 2008) and in pyNEURON (Hines et al., 2009). These tests were run using external HPC and local resources, maximizing available parallel computing. The time resolution for both simulators was 0.1ms. As internal validation tests, some exemplificative ad-hoc structural and functional alternatives (see Fig. 5) were made in the network and then the same simulations were run (in pyNEST). The firing rates of each cell population and their sub-groups affected by stimulus are reported in Table 4 and Fig. 5 to illustrate the similarity of firing rate simulated using pyNEST and pyNEURON.

**Fig. 5.**
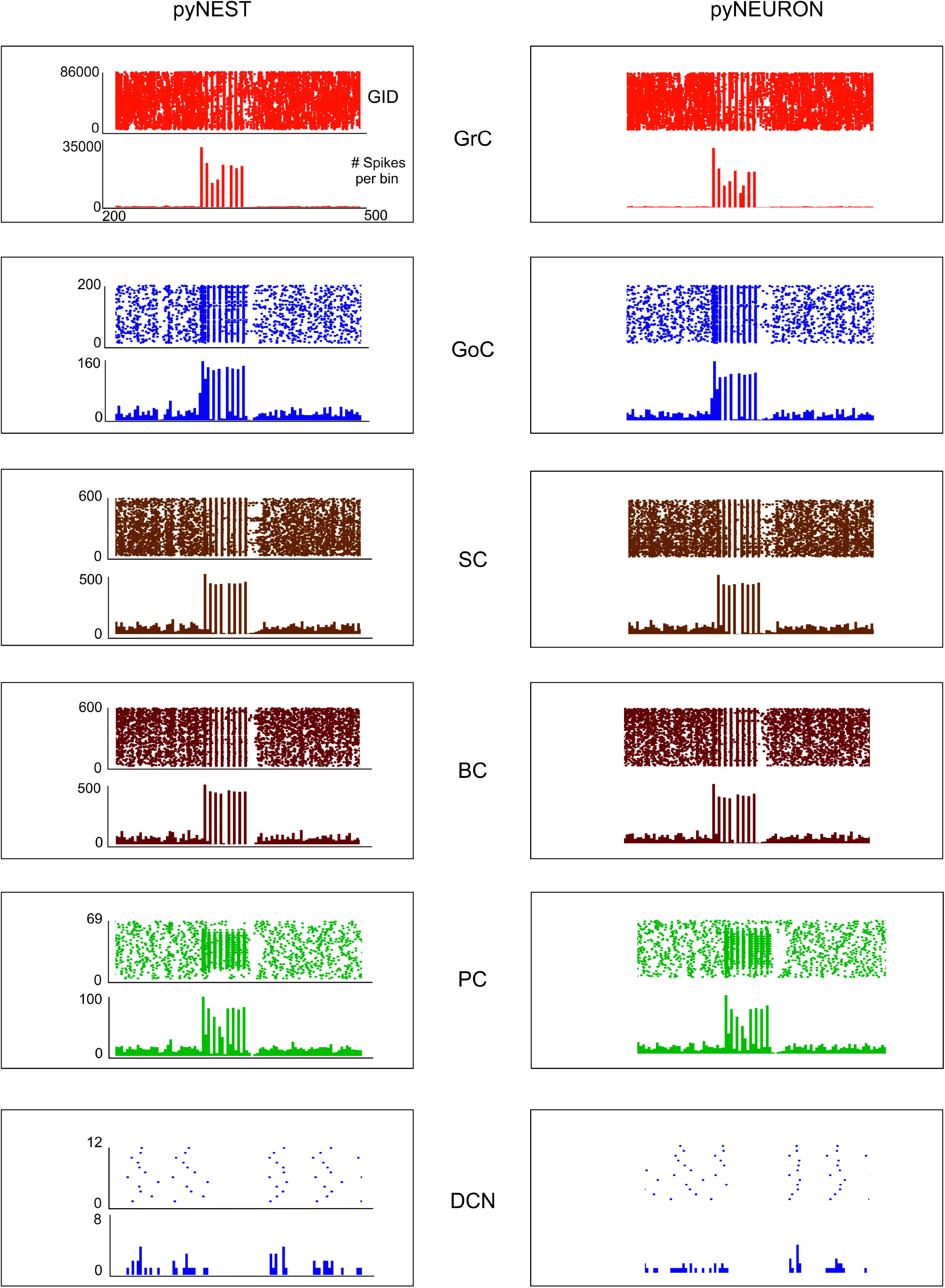
Neuronal discharge. Raster plot and PSTH of the different neuron populations of the cerebellar network model in response to a mossy fiber burst (50 ms at 150 Hz on 2932 gloms) superimposed on a 1 Hz random background. The two simulations used the same cerebellar scaffold and neurons, which were translated from pyNEST into pyNEURON. The basal activity of the different cell populations is visible before and after the stimulus. It should be noted that neuronal discharges are very similar using the two simulators.

**Table 4.** For each neuronal population, the firing rates (mean ± sd) are reported before, during and after stimulation. Excited (inhibited) cells are defined as those increasing (decreasing) the number of spikes during the stimulus (see Methods). Simulation results are shown for pyNEST (white rows) and for pyNEURON (grey rows). In the column “During stim”, the values indicate the firing rates only averaged on the sub-group “Excited (Inhibited) cells”.

| Neuron type | N.cells | N. Excited (or Inhibited) cells (%) | Before (300 ms) [Hz] | During (50 ms) [Hz] | After (300 ms) [Hz] |
| --- | --- | --- | --- | --- | --- |
| Gloms | 7070 | E<sub>NEST</sub>: 2932 (41 %) | 1.0 ± 1.8 | 140.8 ± 4.2 | 0.9 ± 1.8 |
| | | E<sub>NEURON</sub>: 2932 (41 %) | 1.0 ± 1.8 | 140.8 ± 4.2 | 0.9 ± 1.8 |
| GrCs | 88158 | E<sub>NEST</sub>: 26195 (29 %) | 2.0 ± 2.6 | 114.0 ± 32.2 | 1.8 ± 2.5 |
| | | E<sub>NEURON</sub>: 25240(28%) | 2.0 ± 2.6 | 114.8 ± 32.8 | 1.8 ± 2.5 |
| GoCs | 219 | E<sub>NEST</sub>: 146 (66%) | 22.7 ± 13.1 | 157.1 ± 37.2 | 23.5 ± 11.3 |
| | | E<sub>NEURON</sub>: 153 (69 %) | 22.1 ± 12.4 | 154.2 ± 28.9 | 24.9 ± 12.0 |
| SCs | 603 | E<sub>NEST</sub>: 445 (73%) | 33.9 ± 15.7 | 126.2 ± 17.4 | 37.0 ± 14.3 |
| | | E<sub>NEURON</sub>: 452 (74%) | 34.2 ± 15.9 | 131.8 ± 19.6 | 36.9 ± 15.0 |
| BCs | 603 | E<sub>NEST</sub>: 429 (71%) | 30.1 ± 15.1 | 124.1 ± 18.4 | 33.6 ± 14.0 |
| | | E<sub>NEURON</sub>: 447 (74%) | 29.1 ± 15.3 | 123.3 ± 24.9 | 33.8 ± 14.4 |
| PCs | 69 | E<sub>NEST</sub>: 45 (65%) | 58.5 ± 8.5 | 255.5 ± 63.0 | 62.8 ± 8.3 |
| | | E<sub>NEURON</sub>: 47 (68%) | 60.0 ± 9.3 | 256.5 ± 63.8 | 63.0 ± 11.6 |
| DCNCs | 12 | I<sub>NEST</sub>: 12 (100%) | 16.1 ± 1.2 | 0.0 ± 0.0 | 16.3 ± 0.9 |
| | | I<sub>NEURON</sub>: 12 (100%) | 16.6 ± 0.0 | 0.0 ± 0.0 | 16.6 ± 0.0 |

The entire scaffold can be built and run as a Jupyter Notebook in the Brain Simulation Platform (BSP), one of the platforms of Human Brain Project (Markram, 2012). The BSP is an internet-accessible collaborative platform that comprises a suite of software tools and workflows to reconstruct and simulate multi-level models of the brain at different levels of description, from abstract to highly detailed. Here, cells, network and volume configuration parameters can be easily read and modified, since they are stored in a single Python script. Such flexible parametric approach allows to continuously include and tune relevant neurophysiological information and to operate at different simplification levels. A test version of the scaffold model is running on the Brain Simulation Platform at https://www.humanbrainproject.eu/en/brain-simulation/brain-simulation-platform/.

## 3 Results

The cerebellar network is unique for its precise geometrical organization (Fig. 1), which was reconstructed generating a *scaffold model* capable of handling neuronal placement, connectivity and simulations. The neurons were represented as single-point leaky integrate-and-fire (LIF) models(Maas, 1997), tuned to match the input resistance and capacitance, basal discharge and input-output relationship of the specific cerebellar neuron types. The choice of LIF neuron models was motivated by the need to focus first onto the two main construction operations of the scaffold, cell placement and connectivity, and on the role of these latter in determining network properties.

The scaffold is demonstrated here through the exemplar reconstruction and testing of a cerebellar volume of 0.077 mm^3^. The cerebellar cortex volume had 400×400μm^2^ base and 330 μm height subdivided into different layers: molecular layer (150 μm), Purkinje cell layer (30 μm), granular layer (150 μm). The DCN layer had 200×200μm^2^ base (1/4 of cortex) and 600 μm height. As a whole, the model contained96734 cells. It should be noted that these parameters were all user-defined and may be modified depending on the needs, as the model is scalable.

### 3.1 Cell Placement

The *Bounded Self-Avoiding Random Walk* algorithm (see Methods) successfully placed the neurons into all cerebellar regions with the only exception of PCs, which were positioned using a specific algorithm designed to respect their regular spatial organization (Fig. 2A). Fig.2B shows a row of almost equally distanced PCs connected to incoming parallel fibers, faithfully reproducing the typical PCs geometrical organization. These examples show that the placement algorithms can be flexibly configured to account for complex and variable rules of cellular positioning.

As an internal validation, the distribution of pair-wise distances for each cell type was calculated (Fig.2C). For all cell types (except PCs), pair-wise distances were distributed almost normally and the minimum inter-soma distance equated twice the soma radius. As expected, KDE for GrC, GoC, SC and BC pair-wise distances returned a single maximum (at 180.1 μm, 191.0 μm, 184.6 μm and 188.5 μm respectively), while for PCs three local maxima occurred (at 48.6μm, 142.1 μm and 267.0 μm) (for DCNC, KDE analysis was meaningless, given the low number of cells).

### 3.2 Cell Connectivity

The connection rules adopted in this work were designed to account for the rich and specific information available from literature(Eccles et al., 1967; Palay and Chan-Palay, 1974; Korbo et al., 1993), which accounts for convergence/divergence ratios, number of synapses and spatial distribution of axons and dendrites (Fig. 3). The connecting algorithm imposed these geometrical constraints allowing to wire the different neuronal types for a whole of 16 connection types. Five connection types did not require other than these geometric constraints, while pruning was needed in the other 11 cases (either for convergence or divergence or both). The resulting connectome was then compared to the experimental one for validation. Fig.4 shows that the connection ratios of the scaffold were indeed correctly scaling to the physiological ones. Some specific cases are considered below.

Concerning Glom-GrC connectivity, experimental data demonstrated that granule cell dendrites have a maximum length of 40 μm, with a mean value of ~13 μm (Solinas et al., 2010). By imposing a convergence value of 4(each GrC received one Glom on each of its 4-5 dendrites), a mean dendrite length of about ~12μm was found, therefore matching experimental and theoretical determinations (Hamori and Somogyi, 1983; Billings et al., 2014).

Concerning connectivity between the aa and PC dendrites (aa-PC), connections were possible only when the aa-segment was very close to the PC dendritic plane. By analyzing the placement of GrCs in the x-z plane and the vertical extension of the aa, it is estimated that only ~20% of GrCs developed an aa that is sufficiently close to a PC dendrite tree to form a synaptic contact (Bower and Woolston, 1983; Gundappa-Sulur et al., 1999). This estimate was indeed closely matched by the scaffold reconstruction.

Concerning connectivity of parallel fibers with receiving neurons (pf-GoC, pf-SC, pf-BC, pf-PC synapses), the literature is incomplete and shows variable estimates. This most likely reflects difficulties in estimating exact numbers, since the pf can be several millimeters long and they are often cut on the parasagittal plane in histological preparations. In the scaffold reconstruction, the maximum pf length (along z-direction) was bounded to 400 μm (Barbour, 1993; Huang et al., 2006) and pf from GrCs beyond this length were not taken into account.

The statistical distribution of distances between connected cells (Table 4) shows a good matching with anatomical values. This validation of the connectome supports the appropriateness of cell placement and connecting rules.

### 3.3 Neuronal activations in the cerebellar network following mossy fiber stimulation

The aim of these simulations was to assess the emergence of typical spatio-temporal patterns of cerebellar network activity as a consequence of mossy fiber inputs. Simulations were carried out both in pyNEST and pyNEURON showing very similar results. For simplicity, the following data and figures are taken from pyNEST simulations, except for a comparison of the two in Fig.7 and Table 4.

Evoked activity simulating the effect of natural sensory stimulation (Chadderton et al., 2004; Roggeri et al., 2008; Ramakrishnan et al., 2016)was elicited over a noisy background (see above) by a 150Hz – 50msmossy fiber burst. The mossy fiber activity spread over about 0.012 μm^3^ of the granular layer involving 2932 glomeruli out of the 7070 placed in the reconstructed volume. Glomeruli had mean firing rate of~1 Hz before the burst, 140Hz during the burst, and ~1Hz after the burst. The burst induced transient activity changes, specific for each neuronal population, that reverted back to baseline after the end of the stimulus (Fig.7). The sequence of neuronal activations depended on synaptic delays that were set according to physiological data (Eccles et al., 1967)(Fig. 6). The response of the individual neuronal populations was as follows (Fig. 5–6 and Table 4):

- The GrCs discharged at 1.8 Hz at rest and at 114 Hz during burst stimulation, consistent with in-vivo data showing that GrCs had sparse activity characterized by low background firing rates (partly due to the presence of tonic GABAergic inhibition) and high-frequency bursts in response to evoked sensory stimulation (Chadderton et al., 2004).
- GoCs discharged above 22 Hz at rest and above150 Hz during burst stimulation, consistent with in-vivo data (Heine et al., 2010). The basal GoCs firing rate was raised by the noisy background over the autorhythmic frequency and showed a high variability among cells.
- Molecular layer interneurons, SCs and BCs (N=603 for each cell type), discharged at ~30 Hz at rest and above 120 Hz during burst stimulation, consistent with the observation of high-frequency activity during sensory stimulation(Chu et al., 2012).
- PCs discharged at ~58-60 Hz at rest and at ~255 Hz during burst stimulation consistent with in-vivo data (Heine et al., 2010). Interestingly, PCs showed either bursts, or pauses, or burst-pause responses as observed *in* vivo(Herzfeld et al., 2015): out of 69 PCs, the burst was observed in 48 PCs and the pause in 41 PCs. Of the PCs that showed a pause, in 17 PCs it occurred after a burst, while in the other 24 PCs it happened alone.
- DCNCs discharged at ~16 Hz at rest and were completely silenced during burst stimulation. This behavior was expected from the convergent inhibition coming from PCs, supporting the hypothesis that cortico-nuclear synapses act as simplified inverters (Person and Raman, 2012).

**Fig. 6.**
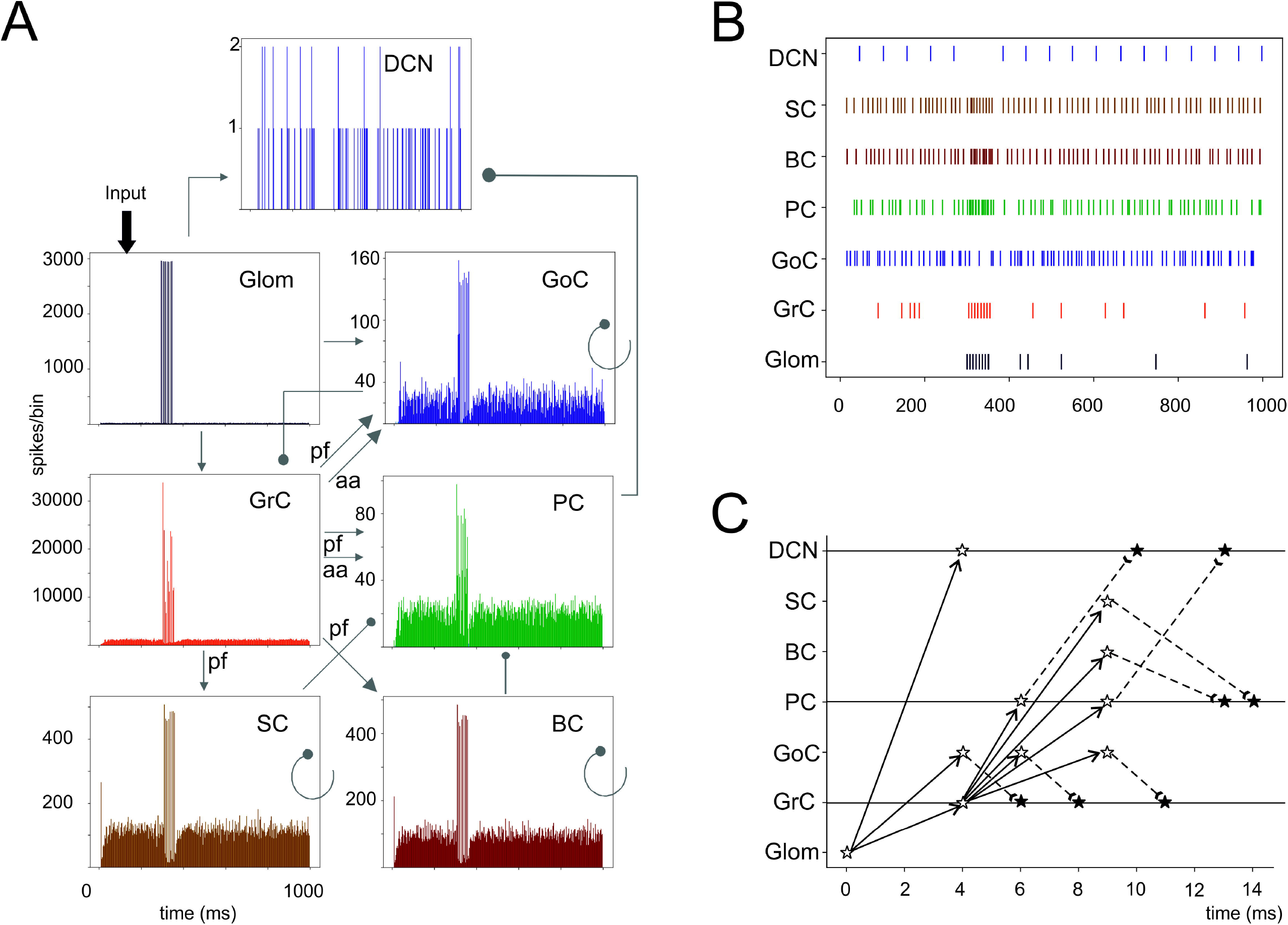
Cerebellar network response to a mossy fiber burst. (**A**) Spikegrams of exemplar cerebellar neurons in the model. A burst in gloms causes a burst-to-burst propagation in GrCs and PCs. GoCs, SCs and BCs also generate bursts that, by being inhibitory, contribute to terminate the GrC and PC bursts and to generate the burst-pause PC response. The DCN cells show a pause during stimulation. (**B**) Raster plot of one cerebellar neuron for each population in the model. Note the spread of the mf bursts inside the cerebellar cortical networks and the corresponding pause in the DCN. (**C**) Spike-time response plot showing the temporal sequence of neuronal activation and inhibition. The arrows represent the connectivity (solid lines show excitatory connections, dashed lines inhibitory connections). The stars represent the post-synaptic neuron response: white stars are excited neurons, black stars are inhibited neurons.

**Fig. 7.**
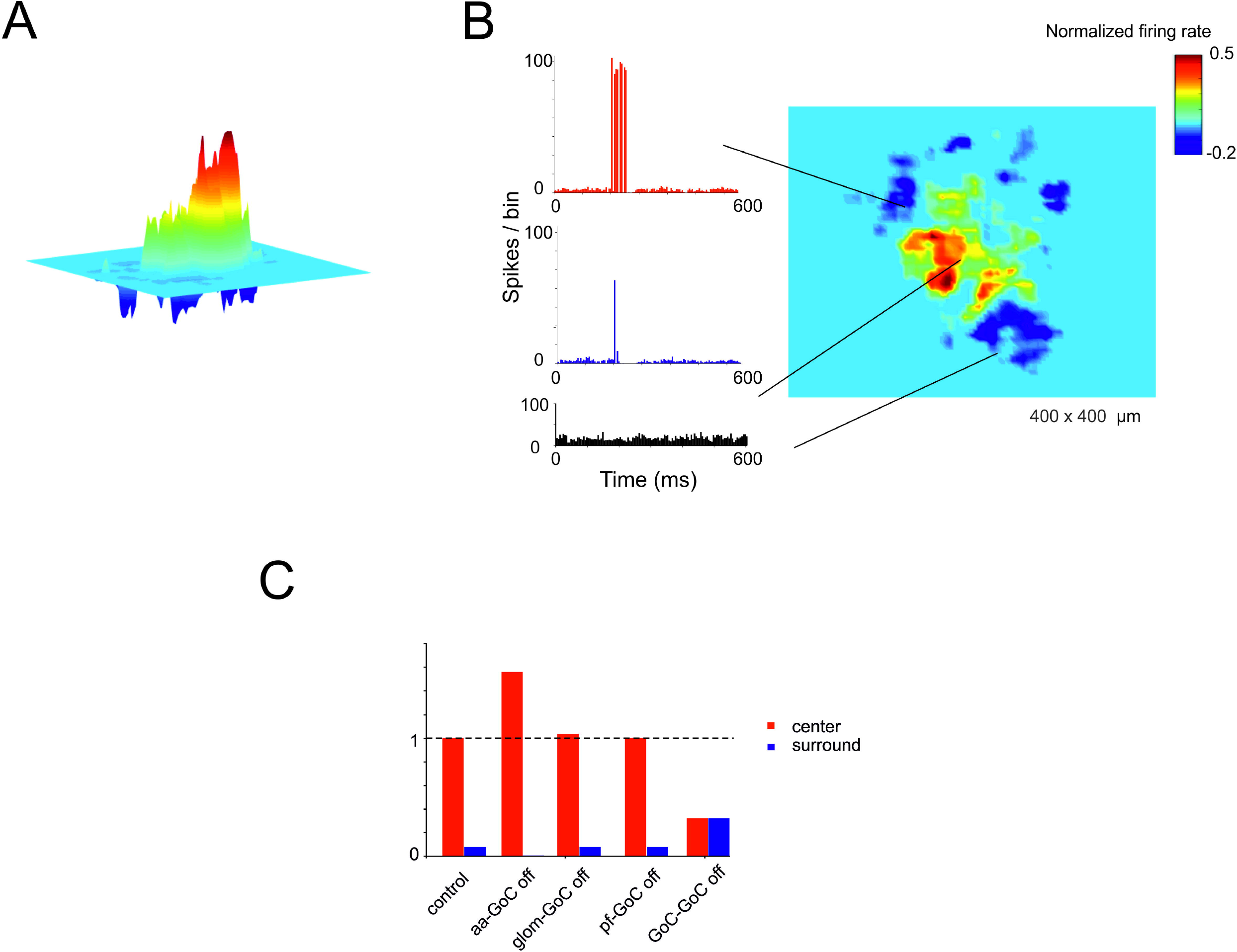
Center-surround organization of activity in the granular layer. (**A**) In response to a mossy fiber burst (40 glom at 150 Hz for 50ms), the granular layer responds with a core of activity surrounded by inhibition. (**B**) PSTH of GrCs in the center-surround. The activity in the core is characterized by robust spike bursts, while just sporadic spikes are generated in the surround. No activity changes are observed outside the center-surround structure. (**C**) The histogram shows the changes in center-surround extension that occur following selective switch-off of synapses impinging on GoCs. Note the prominent role of aa-GoC synapses and GoC-GoC synapses (bars are values normalized to control).

### 3.4 Center-surround organization of granular layer responses

A relevant aspect of network activation that emerged in electrophysiological and imaging experiments is the center-surround organization (Mapelli and D’Angelo, 2007; Diwakar et al., 2011; Gandolfi et al., 2014). In the scaffold, the neuronal response of the granular layer to mossy fiber stimulation showed a typical center-surround organization (Fig.6). This reflected the excitatory/inhibitory ratio (see Methods) with the center more excited than the surround due to lateral inhibition provided by GoCs. The center-surround had a diameter of about 50 μm and GrCs inside the core discharged up to 3-4 spikes organized in a short burst, reflecting previous experimental estimates (Gandolfi et al., 2014). Therefore, the scaffold correctly predicts the consequences of activity in bundles of mossy fibers.

Recently the connectivity of GoCs and GrCs has been extended by the demonstration of new synapses, in particular those between the GrC ascending axon and GoCs (aa-GoC, excitatory)(Cesana et al., 2013) and between GoCs (GoC-GoC, inhibitory)(Hull and Regehr, 2012). The selective switch-off of aa-GoC connections enhanced the center and reduced the surround, the switch-off of GoC-GoC connections reduced the center and increased the surround, while smaller effects followed the switch-off of pf-GoC or mf-GoC synapses (Fig. 9C).

### 3.5 The impact of molecular layer interneurons on PC activation

The molecular layer is critical to regulate PC activity in a way that is still debated (e.g. see(Rokni et al., 2007; Santamaria et al., 2007)).The first assumption is a differential orientation of SC cell axons (mostly transversal or “on-beam”) vs. BC axons (mostly sagittal or “off-beam”) (Eccles et al., 1967). Moreover, both aa and pf are used to activate PCs, as reported in literature (Jaeger and Bower, 1994; Canepari et al., 2001) (Fig. 8). Consistently, in the scaffold model, PC responses were circumscribed into a central spot overlaying the center/surround generated in the granular layer with little diffusion along either transversal or sagittal axis. On both axes, in turn, some PCs were clearly inhibited by the molecular layer interneuron inhibitory network.

**Fig. 8.**
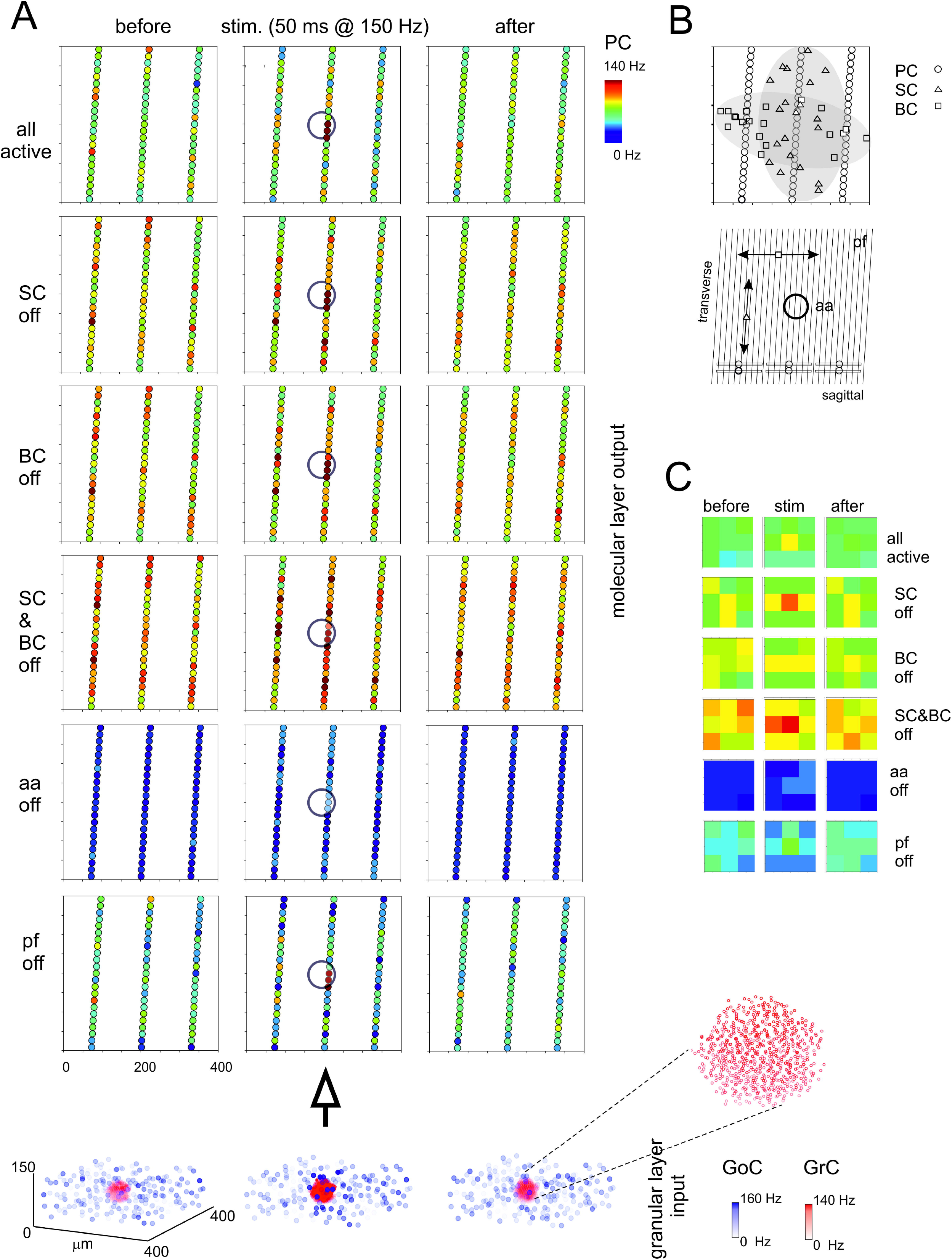
Maps of PC activation and sensitivity to molecular layer connectivity. (**A**) The maps show the activity change of PCs in response to a mossy fiber burst (40 glom at 150 Hz for 50 ms). The pattern of activity is determined by various connection properties that are tested in turn. (**all active**) PC inhibition is achieved through a differential orientation of SC axons (mostly transversal or “on-beam”) *vs*. BC axons (mostly sagittal or “off-beam”) and that PC excitation depends on both aa and pf synapses with specific origin from GrCs. Alternative patterns are generated by (**SC off**) the specific switch-off of SC, (**BC off**) the specific switch-off of BC, or (**SC&BC off**) the complete switch-off of both SC and BC, (**aa off**) the specific switch-off of aa synapses, (**pf off**) the specific switch-off of pf synapses. It should be noted that these changes in network connectivity modify the PC discharge patterns both on-beam and off-beam and extend to a distance that reflects the propagation of activity through the molecular layer interneuron network. The circles indicate the location of the underlying active spots of activity in the granular layer. The bottom plot represents the activity of **GoCs** (blue) and **GrCs** (red) before, during and after the stimulus burst. This activity occurs in a spot (enlarged in the inset) corresponding to the center-surround shown in Fig. 7. (**B**) The schematic diagrams show the orientation of fibers and connections in the network. (**C**) The PC activity was averaged into 3×3 matrices in order to better appreciate where activity changes take place. Note the emergence of the central spot in several cases.

Then, the effect of disconnecting different network elements was tested. Following the switch-off of both SC and BC inhibition, the responsiveness of PCs increased, as expected from SC and BC inhibitory action on PCs. As expected from anatomy, when only BCs were present (i.e. selective switch-off of SCs), excitation extended more effectively along the transverse axis, while when only SCs were present (i.e. selective switch-off of BCs) excitation extended more effectively along the sagittal axis. However, in both cases there was a diffused (though slight) increase of excitation, due to the reduced background inhibition exerted by intrinsic SC and BC discharge. It should also be noted that the activation of PCs in the central spot remained poorly altered, suggesting that these PCs were already nearly maximally activated in control. The selective switch-off of aa synapses caused a diffuse reduction of PC activation, while the selective switch-off of pf synapses had a much smaller effect. Therefore, changes in molecular layer connectivity consistently modified the PC discharge patterns both on-beam and off-beam and extended to a distance that reflects the propagation of activity through the pfs and the molecular layer interneuron network.

### 3.6 Synchronous oscillations caused by noisy background activity in mossy fibers

Recordings from the granular layer *in vivo* have revealed low-frequency local field potential oscillations that occur synchronously over distances of several hinders of micrometers (Pellerin and Lamarre, 1997; Hartmann and Bower, 1998). Similar properties were observed also in previous granular layer models (Maex and De Schutter, 1998; Solinas et al., 2010; Sudhakar et al., 2017). In the scaffold model, spontaneous circuit activity clearly emerged due to background firing in the mossy fibers, provided that the frequency of the background mossy fiber discharge was increased from 1 Hz to 5 Hz and pfs-GoCs connection weight was increased from 0.2 to 30.4, supporting the concept that oscillations require a specific synaptic balances to emerge (Maex and De Schutter, 1998; Solinas et al., 2010; Sudhakar et al., 2017) (Fig. 9). In response to the input, GrCs sparsely discharged at low frequencies (GrCs do not show intrinsic spontaneous activity), while the intrinsic activity of all the other neurons was modulated (GoC, PC, MLI and DCNC are autorhythmic) (see *I_e_* values in Table 3). Interestingly, the neurons of the granular layer (GrCs and GoCs) showed a pattern of low-frequency oscillations (mean frequency of 1.8 Hz) that was evident across the whole network. The oscillation frequency is the same in autocorrelograms of both GrCs and GoCs, and in the cross-correlogram between Golgi and granule cells. This ensemble behavior is probably due to the inhibitory feedback from GoCs to GrCs in the following way: (1) GrCs activity sums up in several GoCs, (2) GoCs, which are synchronized through parallel fibers and reciprocal inhibitory synapses, discharge almost synchronously, (3) a large population of GrCs is phasically inhibited, (4) inhibition terminates and GrCs recover responsiveness to the background mossy fiber input, then restarting the cycle.

**Fig. 9.**
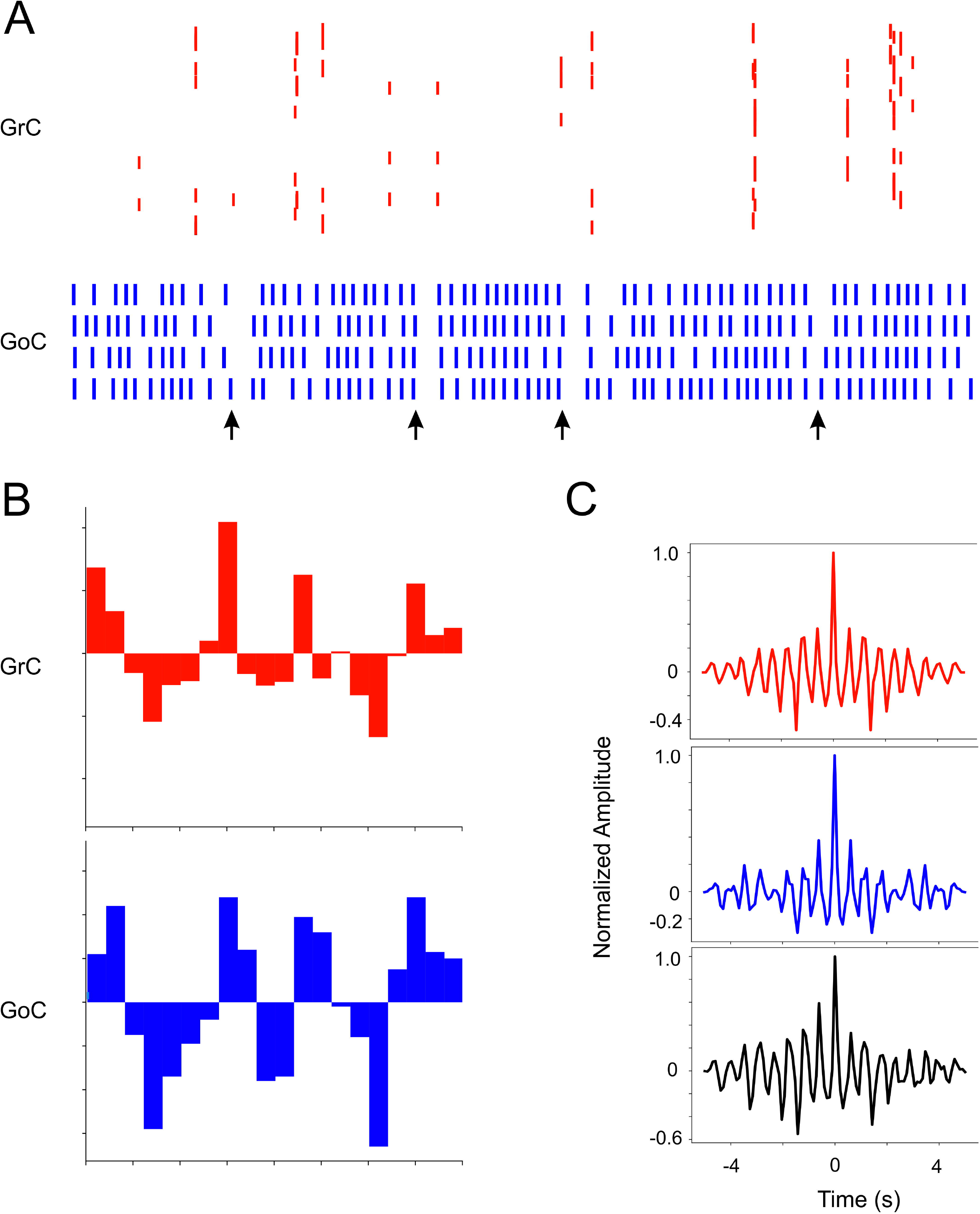
Coherent low-frequency oscillations in granular layer neurons. Activity of GrCs (red) and GoCs (blue) during sustained 5 Hz random mf input. (**A**) Raster plots from exemplar GrCs and GoCs. Note that synchronous patterns are visible in the neuronal response (arrows). In this regimen, GoC activity is more intense than GrC activity due to the autorhytmic discharge of GoC neurons. The neurons are not necessarily part of a center-surround and therefore not all activities appear correlated. (**B**) Cumulative PSTH of the whole GrC and GoC populations of the model along a 5 sec period. Note that the two PSTH show marked low-frequency oscillations (average 1.8 Hz) around their average level of activity. (**C**) Autocorrelograms of activity in the GrC and GoC populations and crosscorrelogram of the GrC and GoC populations (in this example the inhibition among GoCs is switched off). Note the high level of correlation in all the three cases on the same main frequency of1.8 Hz.

## 4 Discussion

In this paper a new scaffold modeling strategy is presented, that is used to simulate fundamental functional properties of the cerebellar microcircuit. The cerebellar scaffold includes the canonical neuron types (GrCs, GoCs, PCs, SCs, BCs, DCNCs), each one with specific geometry of dendritic and axonal fields and with specific convergence/divergence ratios for connectivity. The single neuron models were purposefully simplified in order to focus on network connectivity properties. The circuit functionality was then tested by applying background activity and burst stimuli and evaluating the network responses. In addition to faithfully reproduce a broad range of experimental observations, the cerebellar scaffold shows the emergence of complex spatiotemporal patterns of activity similar to those observed *in vivo* and eventually predicts the critical role of local connectome for network functionality.

### 4.1 The scaffold design

The scaffold includes two modules: cell placement and connectivity. The first module placed neurons in their corresponding layers according to density values derived from literature. Cell placement exploited a bounded self-avoiding random walk algorithm, except for PCs, which required a placement rule accounting for their regular disposition and quasi-planar non-intersecting dendritic trees. The second module generated microcircuit connectivity by defining the pre- and post-synaptic neurons among those intersecting their dendritic and axonal fields and then establishing the corresponding number of synapses through specific connection probabilities. Geometrical constraints and divergence/convergence ratios derived from literature played a critical role to implement the microcircuit connectome. The distributions of soma distances, both for cell positioning and connectivity, were assessed and provided an internal validation for the network construction processes. The cerebellar scaffold was then implemented using LIF neuron models, whose parameters were tuned to approximate the basal firing and input-output relationships of cerebellar neurons. Finally, functional simulations required the scaffold to be uploaded into a neuro-simulator, either pyNEST or pyNEURON, that worked equivalently well for this purpose.

The scaffold modeling strategy used for the cerebellum microcircuit differs from that used for the cortical microcolumn mostly because here the connectivity rules are based on available statistical and geometrical information rather than on single neuron morphologies and touch-detection (Markram et al., 2015). This allows the scaffold to fully exploit the experimental data available in the cerebellar literature despite incomplete availability of detailed morphological reconstructions of cerebellar neurons. By considering that neuron models based on detailed morphological reconstructions are still unavailable in most neuronal circuits, the strategy adopted here for the cerebellum has a large potential for applicability in a variety of different brain microcircuits. It should be noted that our “intersection-connection” rule is formally similar to the “proximity-connection” rule used for touch-detection in (Markram et al., 2015). Eventually, the touch detection strategy could be implemented in the scaffold providing a construction alternative, in which connectivity is directly constrained by neuronal morphology. The advantage would be to specifically connect synapses on specific positions of the dendritic tree, fully exploiting non-linear dendritic computations. Also the data positioning rules could be changed, for example by importing cell positions from the Allen Brain Atlas directly (available at https://portal.bluebrain.epfl.ch/) or using network growing algorithms (Setty et al., 2011; Nguyen et al., 2016).

### 4.2 Simulation and Validation of Cerebellar Network Properties

As a consequence of background input firing and of punctuate sensory stimulation (D’Angelo et al., 2016), the scaffold model predicts a set of relevant network response properties that match experimental observations. These include:

- Loose synchronicity in GrC and GoC activity in response to random mossy fiber inputs (Pellerin and Lamarre, 1997; Hartmann and Bower, 1998). Similar properties were observed also in previous granular layer models (Maex and De Schutter, 1998; Solinas et al., 2010; Sudhakar et al., 2017).
- Center-surround organization of GrC discharge in response to activation in mossy fiber bundles, that was reported *in vitro* and predicted also to occur *in vivo* (Mapelli and D’Angelo, 2007; Diwakar et al., 2011; Gandolfi et al., 2014).
- Combination of bursts and pauses in PC and DCNC responses following mossy fiber burst stimulations (Herzfeld et al., 2015).It should be noted that these combinations of bursts and pauses occurred purely on a network connectivity basis, suggesting that additional single-neuron mechanisms would reinforce the mechanism (Masoli et al., 2015; Masoli and D’Angelo, 2017).
- Regulation of the spatial PC discharge pattern depending on the geometry of SC and BC inhibition, predictive of molecular layer patterns of activity *in vivo* (Santamaria et al., 2007).
- Enhancement of local PC responses by aa (Bower and Woolston, 1983; Walter et al., 2009; Cesana et al., 2013). The emergence of “spots” in PC activity is still debated (Rokni et al., 2007; Santamaria et al., 2007) but these simulations suggest that the vertical organization of GrC - PC transmission based on aa provides indeed a robust structural substrate for spots, as they persisted in most of the synaptic configurations that were simulated.
- Enhancement of center-surround responses due to the aa (Cesana et al., 2013). This observation suggests that the GrC aa has a local effect on granular layer as well as on molecular layer activity.

Interestingly, despite the use of simplified LIF neuron models, the observation of these activity patterns in the model suggests that structural constraints play a critical role in determining local neuronal dynamics. There are several aspects that remain to be assessed and will be easily incorporated into more advanced versions of the cerebellar scaffold.

First of all, assessing the role of non-linear neuronal properties, like intrinsic oscillations, resonance bursting and rebounds, requires to incorporate into the scaffold realistic ionic-channel based neuronal models. Along with this, dendritic computation needs morphologically detailed neuron models that are currently under construction and testing. These include the PCs model (e.g. (Masoli et al., 2015; Masoli and D’Angelo, 2017)), the GrC model (Masoli et al., 2017), the GoC model (Solinas et al., 2007a, b), the SC and BC model (currently under construction), the DCN model (Steuber and Jaeger, 2013). Dynamic synapses (Tsodyks and Markram, 1997; Nieus et al., 2006; Migliore et al., 2015) will likewise be incorporated replacing the current exponential representations.

Secondly, the scaffold could be used to evaluate the trade-off between computational efficiency and precision. Therefore, the present LIF single point neurons could be substituted by others (E-GLIF) embedding non-linear firing properties (e.g. (Geminiani et al., 2018)) and accounting for synaptic dendritic location by modifying the transmission weight depending on the distance of synapses from the soma (Rössert et al., 2016).

Thirdly, fully implementing cerebellar connectivity requires the introduction of models of the inferior cerebellar olive (IO) (Libster and Yarom, 2013; De Gruijl et al., 2014). This will complete the DCN-PC-IO cerebellar circuit, allowing the model to simulate oscillations in the olivo-cerebellar circuit, their impact on PC dendritic calcium signaling and computation, and eventually climbing fiber control of plasticity at parallel fiber synapses (Coesmans et al., 2004).

Finally, the addition of novel connections and cells, like the PC to GrC inhibition (Guo et al., 2016) in the anterior cerebellum, the unipolar-brush cell subcircuit in the flocculo-nodular lobe (Mugnaini and Floris, 1994; Subramaniyam et al., 2014), or the DCN to granular layer connections (Gao et al., 2016) will allow to further expand the simulation of cerebellar processing in different cerebellar modules.

## 5 Conclusions

The scaffold model was able to reconstruct the complex geometry and neuronal interactions of the cerebellar microcircuit based on *intersection-connection* rules. Given its architectural design, that puts in series interchangeable programming modules, the scaffold could now be used to plug-in different network configurations into neuronal simulators like e.g. pyNEST and pyNEURON (Brette et al., 2007; Hines et al., 2009). Both the cell placement algorithm, the neuron model types and the connectivity rules could be substituted to assess different construction strategies and adapt to available data to probe specific functional hypotheses. For example, the connectivity could be recalculated using realistic neuronal morphologies and touch-detection algorithms (*proximity-connection* rule), as in the cortical microcolumn model (Markram et al., 2015). We envisage that this scaffold modeling strategy, given its versatility, will be also able to host microcircuits different from cerebellum providing a new tool for neurocomputational investigations. Eventually, translation of the scaffold model into PyNN would also facilitate neurorobotic and neuromorphic hardware applications (Davison et al., 2008).

## 7 Author Contributions

SC, EM, CM and CC have written the code and run the simulations and taken part to the production of text and figures. CC has coordinated the modeling and simulation activity. ED has directed the work and edited the manuscript.

## 9 Acknowledgments

This research was supported by the HBP Neuroinformatics Platform, HBP Brain Simulation Platform, HBP HPAC Platform, funded by the European Union’s Horizon 2020 Framework Programme for Research and Innovation under the Specific Grant Agreement No. 785907 (Human Brain Project SGA2).

